# *Phosphate transporter* (*Pht*) gene families in rye (*Secale cereale* L.) – genome-wide identification and sequence diversity assessment

**DOI:** 10.1101/2024.08.09.607312

**Authors:** David Chan-Rodriguez, Brian Wakimwayi Koboyi, Sirine Werghi, Bradley J. Till, Julia Maksymiuk, Fatemeh Shoormij, Abuya Hilderlith, Anna Hawliczek, Maksymilian Królik, Hanna Bolibok-Brągoszewska

**Author notes:** Corresponding author: Hanna Bolibok-Brągoszewska.

## Abstract

**Background:** Phosphorus is a macronutrient indispensable for plant growth and development. Plants utilize specialized transporters (PHT) to take up inorganic phosphorus and distribute it throughout the plant. The PHT transporters are divided into five families: PHT1 to PHT5. Each PHT family has a particular physiological and cellular function. Rye (*Secale cereale L*.) is a member of *Triticeae*, and an important source of variation for wheat breeding. It is considered to have the highest tolerance of nutrient deficiency, among *Triticeae*. To date, there is no report about genes involved in response to phosphorus deficiency in rye. The aim of this study was to: (i) identify and characterize putative members of different phosphate transporter families in rye, (i) assess their sequence diversity in a collection of diverse rye accessions via low-coverage resequencing (DArTreseq), and (iii) evaluate the expression of putative rye *Pht* genes under phosphate-deficient conditions.

**Results:** We identified 29 and 35 putative *Pht* transporter genes in the rye Lo7 and Weining reference genomes, respectively, representing all known *Pht* families. Phylogenetic analysis revealed a close relationship of rye PHT with previously characterized PHT proteins from other species. Quantitative RT PCR carried out on leaf and root samples of Lo7 plants grown in Pi-deficient and control condition demonstrated that *ScPht1;6, ScPht2* and *ScPht3;1* are Pi-deficiency responsive. Based on DArTreseq genotyping of 94 diverse rye accessions we identified 820 polymorphic sites within rye *ScPht*, including 12 variants with a putatively deleterious effect. SNP density varied markedly between *ScPht* genes.

**Conclusions:** This report is the first step toward elucidating the mechanisms of rye’s response to Pi deficiency. Our findings point to multiple layers of adaptation to local environments, ranging from gene copy number variation to differences in level of polymorphism across *Pht* family members. DArTreseq genotyping permits for a quick and cost-effective assessment of polymorphism levels across genes/gene families and supports identification and prioritization of candidates for further studies. Collectively our findings provide the foundation for selecting most promising candidates for further functional characterization.

## BACKGROUND

Plants are sedentary organisms, forced to rely on the supply of nutrients available in their immediate vicinity. During evolution, plants have developed complex and intricate mechanisms of efficient nutrient uptake and utilization. Phosphorus (P) is a macronutrient indispensable for plant growth and development, especially for the development of roots. At the cellular level, various metabolic processes, such as synthesis of nucleic acids, require P [1].

Agricultural soils often impose low-P environments on plants, leading to losses in crop yields [2, 3]. Orthophosphate (Pi) is the phosphorus form bioavailable for plants. The pool of bioavailable P is strongly affected by the soil pH [4]. Current agricultural practices utilize fertilizers to boost crop production in such nutrient-deficient soils. However, the only source of P used for fertilizer production comes from phosphate rock, which is a non-renewable resource, close to depletion. Some countries, such as the USA and China, have stopped exporting phosphate rock for strategic reasons [5]. Modern crop varieties rely on intense Pi fertilizing to maintain high yields. As a consequence, these crop varieties exhibit poor performance in the widely spread low-P agricultural systems [2, 3, 6]. Plants utilize a relatively small portion of applied P fertilizers (up to 30%), because of the soil P fixation – formation of Al and Fe oxides at acidic pH, calcium phosphates at alkaline pH, and biological fixation by soil microbiome – into plant-inaccessible form [4, 7]. Therefore, elucidating the mechanisms of P use efficiency and low P tolerance will facilitate the breeding of new crop varieties to meet the food demands of the growing human population.

Plants require to maintain adequate levels of P within the cells and tissues to develop and function properly. It is a complex task to orchestrate the primary uptake, translocation, and allocation of Pi to deliver it where and when the plant needs it. Thus, P homeostasis machinery must run smoothly to keep the plants from under or over accumulating P. During low P conditions, the expression of genes involved in Pi mobilization increases [8]. These genes collectively are known as phosphate-starvation inducible genes (PSI) The molecular mechanism governing the transcriptional regulation of PSI genes – which include phosphate transporters – has been extensively studied in *Arabidopsis thaliana*. For instance, the transcriptional factor Phosphate starvation response 1 (PHR1) binds the *cis*-regulatory element P1BS (PHR1-binding site) to positively regulate the expression of PSI, which includes some members of the *Pht* gene families [9]. Similarly, Pi deficiency regulates the rice (*Oryza sativa*) PSI genes, suggesting a conserved regulatory mechanism across plant species [10].

Plants utilize specialized transporters to take up inorganic Pi from the rhizosphere and distribute it throughout the plant. The PHT transporters are divided into five families, namely PHT1, PHT2, PHT3, PHT4, and PHT5. Each PHT transporter family has a particular physiological and cellular function in plants [11, 12]. The PHT transporter family sizes vary across the plant species. For example, Arabidopsis and rice differ in the number of PHT1 and PHT3 members. The Arabidopsis PHT1 family includes nine members, while the rice contains 13 [8]. Similarly, the rice PHT3 family has three more members than Arabidopsis PHT3 family; six and three members in rice and Arabidopsis, respectively. The members of PHOSPHATE TRANSPORTER 1 (PHT1) membrane-localized transporter family are associated with the primary uptake, translocation, and allocation of Pi [13, 14]. Some PHT1 members are strongly expressed in vegetative (leaves, stems, and roots) and reproductive tissues (flowers) under Pi deficiency [15]. The other transporter families mobilize Pi to internal compartments such as the chloroplast (PHT2), mitochondria (PHT3), Golgi and plastids (PHT4), and vacuole (PHT5) [13, 14]., The PHT1 transporter family has been extensively studied in crop plants including rice [16], barley [17], maize [18], wheat [19], *Setaria italica* [20], sorghum [21, 22], finger millet [23] and poplar [12]. Unlike the rest of the Pi transporters, the PHT2 family contains a single member and an expression restricted to leaves in most plants with the exception of poplar [12, 24]. PHT3 proteins are located in the mitochondria and mediate the distribution of Pi between the inner mitochondrial membrane and cytoplasm [25, 26]. PHT4 proteins are localized in the chloroplasts, the Golgi apparatus and non-photosynthetic plastids [27]. In Arabidopsis, the PHT5 imports Pi to the vacuole to maintain the Pi homeostasis in the cytoplasm [11].

The natural genetic variation within plant species is crucial for identifying unique alleles or rare genetic variants to develop more resilient and productive plant varieties, especially for the complex trait of low P tolerance. Various efforts have been made to map loci associated with low P tolerance in model plants and crops. Several studies have highlighted the use of genome-wide association studies (GWAS) in successfully identifying natural allelic variations associated with P deficiency tolerance in *Aegilops tauschii* [28], Arabidopsis [29], soybean [30], wheat [31], and maize [32]. For example, a GWAS conducted in 277 Arabidopsis accessions identified three genes controlling root growth in response to low-Pi conditions [33]. Another study evaluating 82 bread wheat accessions identified nine marker-trait associations involving high-confidence candidate genes. Some of those candidate genes are influenced by P deficiency [34]. In rice, a GWAS study identified the *OsAAD* gene contributing to P utilization efficiency and grain yields [35]. The *PSTOL1* gene is an example of the potential of natural variation in crop improvement for P deficiency tolerance in rice. This gene is absent from the P-starvation-sensitive rice variety Nipponbare, but present in the P-deficiency-tolerant variety Kasalath – a traditional rice variety [36].

Rye (*Secale cereale* L.) is a cereal especially popular in Central, Eastern and Northern Europe. Poland is the world’s second largest rye grain producer (http://www.fao.org/faostat). Genetically rye is a diploid, with a very large (8Gpz) and highly repetitive genome [37]. Rye is a member of *Triticeae*, closely related to wheat and barley, and an important source of variation for wheat breeding [38]. Frequently grown on marginal soils, rye is considered to have the highest tolerance of abiotic stresses, such as frost, drought, aluminum toxicity, among *Triticeae*, and the highest tolerance of nutrient deficiency [39, 40]. Contrary to barley and wheat, rye is out-crossing, and has a genetic self-incompatibility mechanism. Therefore, the development of homozygous lines for breeding and research is a daunting and time-consuming task. The rye genetic diversity remains underutilized as a nutrient-deficiency-tolerant allele resource for crop improvement, and there is a lack of knowledge about the genetic basis of P deficiency tolerance in rye.

This study identified and characterized putative members of different phosphate transporter families in *Secale cereale* L. Lo7 [37] and Weining reference [41] genomes. Our study also assessed the sequence diversity of the identified putative rye phosphate transporters (*ScPht*) in a collection of 94 diverse rye accessions via low-coverage resequencing (DArTreseq). Furthermore, we evaluated the gene expression of some of these putative *ScPht* under Pi-deficient conditions.

## METHODS

### Plant material and growth conditions

Tissue for gene expression analysis was collected from plants of rye inbred line Lo7 grown in hydroponics in Pi-deficient and control conditions. Seeds of the inbred line Lo7 were germinated in a growth room in transparent polyethylene containers lined with a thin layer of moist horticultural pumice gravel. After six days, uniformly germinated seedlings without residual endosperm were transferred to hydroponics. Nutrient solution based on [42] was used. The control medium contained 0.2 mM KH_2_PO_4_ and the Pi-deficient medium contained 0.004 mM KH_2_PO_4_. Plants were grown in a growth chamber at 19 °C and photon fluency rate of ca. – 500 µmol m^-2^s^-1^, (14 hours day/10 hours night). Nutrient solution was exchanged three times per week. Leaf and root samples were collected on days 14 and 21 of the hydroponic trial. All tissue samples were frozen in liquid nitrogen and stored at −80°C.

DArTreseq genotyping was carried out on DNA samples of 94 rye accessions. The accessions were selected to cover a broad spectrum of rye genetic variation in terms of geographic origin and improvement status and comprised: 16 inbred lines, five modern cultivars, 16 historic cultivars, 39 landraces, eight wild/weedy accessions and ten accessions of unknown improvement status. Detailed information on the accessions used in this study is provided in the Additional File 1 (Table S1. Each accession was represented by a single plant. Germination of seeds and plant cultivation for leaf tissue collection, as well as DNA isolation using Mag-Bind Plant DNA DS Kit (OMEGA Bio-Tek) was carried out as described in [43].

### Identification of *PHT* transporters in Lo7 and Weining rye genomes

Putative phosphate transporters belonging to the PHT1, PHT2, PHT3, PHT4 and PHT5 families were identified in the inbred line Lo7 and the Chinese elite rye Weining genomes using both the TBLASTN algorithm within the IPK Galaxy Blast Suite (https://galaxy-web.ipk-gatersleben.de/) and in WheatOmics 1.0 web platform (http://wheatomics.sdau.edu.cn), respectively. We used the Arabidopsis and rice PHT transporters - PHT1.1 (NP_199149.1; AAN39042.1), PHT2.1 (NP_189289.2; XP_015626495.1), PHT3.1 (NP_196908.1; NP_001406294.1), PHT4.1 (NP_180526.1; BAS71585.1) and PHT5.1 (NP_001185297.1) - as protein sequence queries. The rye Lo7 and Weining *Pht* coding sequences were retrieved from the IPK Galaxy Blast Suite and WheatOmics, respectively.

### Chromosomal location, gene structure, and amino acid sequence analysis

We obtained information about the chromosomal location and exon-intron structure of each putative rye *Pht* from the Ensembl plant database (https://plants.ensembl.org/Secale_cereale/Info/Index). The predicted amino acid sequences for all putative rye PHT were retrieved using the ORFinder tool from NCBI (https://www.ncbi.nlm.nih.gov/orffinder/) and the Ensembl plants database. We examined the protein sequences for conserved domains using the InterPro (https://www.ebi.ac.uk/interpro/) and SMART (http://smart.embl.de/) tools. We predicted the subcellular localization and the transmembrane helixes for each rye PHT member using the Plant-mPLoc database (http://www.csbio.sjtu.edu.cn/bioinf/plant-multi/), Plant-mSubP (http://bioinfo.usu.edu/Plant-mSubP/) and TMHMM Server v. 2.0 web tool (https://services.healthtech.dtu.dk/services/TMHMM-2.0/). The chromosome diagrams showing the location of the *Pht* genes were generated using the Phenogram web tool (https://visualization.ritchielab.org/phenograms/plot). We generated the gene structure and protein conserved domain diagram using the Gene Structure Display Server (GSDS 2.0) and TBtools, respectively [44, 45].

### Analysis of the *ScPht*s promoter regions

We analyzed the 2 Kb upstream region of the rye *Pht* genes using the plant cis-acting regulatory DNA element database PLACE (www.dna.affrc.go.jp/PLACE/?action=newplace). The upstream region sequences for all putative rye Lo7 *Pht* genes were obtained from the IPK Galaxy Blast Suite (https://galaxy-web.ipk-gatersleben.de/) and the Ensembl plants database (https://plants.ensembl.org/Secale_cereale/Info/Index).

### Phylogenetic analysis

We performed multiple alignments of the PHT protein sequences from *Secale cereale* Lo7 and Weining, *Oryza sativa, Sorghum bicolor*, and *Arabidopsis thaliana* using Clustal W. The Arabidopsis, rice, and sorghum PHTs protein sequences were retrieved from the Phytozome database (https://phytozome-next.jgi.doe.gov)(Additional file 1: Table S2). We maintained the pairwise alignment default parameters (Gap Opening Penalty:10.00; Gap Extension Penalty: 0.10) but adjusted the multiple alignment parameters (Gap Opening Penalty: 3; Gap Extension Penalty: 1.8). To generate the unrooted PHT protein tree, we used the Neighbor-Joining method implemented in MEGA X [46]. The bootstrap values were calculated from 1000 replicates. For the phylogenetic analysis of monocot PHT1, we carried out multiple alignments of PHT1 transporters of *Secale cereale, Triticum aestivum, Hordeum vulgare, Oryza sativa, Sorghum bicolor, Zea mays, Setaria viridis, Brachypodium distachyon*, and *Arabidopsis thaliana*. The PHT1 protein tree was built following the parameters described above. We used Figtree v1.4.4 software (https://github.com/rambaut/figtree) to further annotate and visualize the protein phylogenetic tree.

### Quantitative RT-PCR

Total RNA was extracted using the Universal RNA Purification Kit (EURX, Gdansk, Poland) according to the manufacturer’s instructions. RNA was assessed for quality using a Nabi UV/Vis Nano Spectrophotometer and for integrity on 2% denaturing agarose gel. RNA (∼1 μg) was used to synthesize first-strand cDNA using the RevertAid First Strand cDNA Synthesis kit (Thermo Fisher Scientific, Vilinus, Lituania). Quantitative PCR was performed in a 20 μl reaction mixture containing 10 μl of FastStart Essential DNA Green Master (Roche Diagnostics, Mannheim, Germany), 8μl of 2.5 ng µl-1 cDNA, and 1 μl of 10 mM of each primer using LightCycler® 96 System (Roche Diagnostics, Mannheim, Germany). The thermal cycler was programmed as follows: one cycle at 95°C for 10’, followed by 40 cycles at 95°C for 10 s, 60 °C for 10 s, and 72°C for 15 s, and the melting step - 95°C for 10 s, 65°C for 60 s and 97°C for 1 s. The *ScActine* and *ScEFa1* genes were used as reference genes [47]. The coding sequences of the identified *ScPht* genes were aligned using the Clustal Omega web tool [48]. This alignment facilitated the analysis of the similarity percentage among the sequences, allowing us to identify *ScPht* genes with the least similarity for primer design. Multiple primers specific to the following genes *ScPht1;6, ScPht1;7, ScPht1;11, ScPht2, ScPht3;1, ScPht3;4, ScPht3;5* and *ScPht5;3* were designed using Primer-BLAST [49]. All primers were checked for amplification efficiency and only primers that demonstrated an efficiency of at least 90% were selected for the experiment. Reactions for the reference gene were included in each plate. Each reaction was performed in two technical replicates and three biological replicates, and the data from RT-qPCR were analyzed using the 2–ΔΔCt method [50]. Kruskal test was used to assess statistical significance.

Statistical analyses and plots were done using R [51] packages ggplot2 (v3.3.3) [52] and dplyr (v0.7.6) (https://github.com/tidyverse/dplyr).

### DArTreseq® genotyping - evaluation of sequence diversity of putative *ScPht* genes

Genotyping via DArTreseq® was carried out at Diversity Arrays (Bruce, ACT, Australia). This approach, involving an effective depletion of repetitive sequences, represents a novel way of acquiring a very detailed genome profile in a more cost-effective manner than the Whole Genome Sequencing (WGS) methods. For removal of methylated genomic DNA each sample was digested with *MspJI* (New England Biolabs, Massachusetts, USA). After purification with AMPure beads (Beckman Coulter, California, USA) DNA fragments larger than 300 bp were retained and used for library preparation. Nextera DNA Flex Library Prep kit (Illumina, California, USA) was used for this purpose and each library was provided with an unique dual barcode. Sequencing was performed on a 150 cycles pair-end run on Illumina Novaseq6000 sequencer (California, USA) to achieve 10 to 20 million reads per sample. Raw sequence reads were processed in DArTreseq pipeline. Poor quality reads were discarded and remaining sequences aligned to Lo7 genome sequence [37]. FreeBayes (https://github.com/freebayes/freebayes) was used to call SNP markers. Genetic variants were annotated for their predicted effect on gene function using SIFT [53]. A SIFT database was created using the Lo7 genome reference, the associated GFF3 annotation file converted to GTF using gffread, and the UniRef90 protein database using SIFT4G_Create_Genomic_DB (Pertea end Pertea 2020). The effect of missense changes on protein function were predicted using the SIFT4G annotator tool [53]. For evaluation of genetic diversity of 94 rye accessions SNPs with MAF > 0.01 and <10% missing data were identified. Then, an Euclidean distance matrix was calculated using R package ’stats’ [51] and used for NJ tree construction in MEGA11 [54] and principal coordinates analysis in GenAlEx 6.5 [55, 56].

## RESULTS

### Identification of *Pht* family members in rye

We identified 29 and 35 putative *Pht* transporter genes in the rye Lo7 and Weining reference genomes, respectively (Additional File 1: Table S3 and Table S4). Both reference genomes possess 1 *Pht2*, 2 *Pht4* and 4 *Pht5* members. However, the Lo7 and Weining genomes differ in the number of members of the *Pht1* and *Pht3* families. The rye Lo7 genome contains 16 *Pht1* and 6 *Pht3* members, while the rye Weining genome holds 19 *Pht1* and 9 *Pht3* putative transporter genes (Figure 1). We designated names for rye *Pht* genes (*ScPht*) according to the phylogenetic relationship to the Arabidopsis and rice PHT transporters. We verified that every putative PHT transporter contains the characteristic conserved domain of each PHT family. Tables S3 and S4 (Additional File 1) show the gene ID, the size of genomic loci, coding sequence and amino acid sequence, as well as the conserved domain, subcellular localization, and the transmembrane helices for each putative rye PHT member from the Lo7 and Weining genome, respectively. Figure 1A illustrates that in the Lo7 genome the *ScPht* genes are distributed across all chromosomes, except for the chromosome 1R. Most *ScPht* genes are located on chromosome 7R, with ten *ScPht1* genes and one *ScPht*3 gene. Chromosomes 6R and 5R contain five *ScPht* genes each. Chromosome 4R contains three *ScPht* genes, while the chromosomes 3R and 2R contain only two *ScPht* genes each. In the Weining genome the *Pht* genes are scattered across six chromosomes, too (Figure 1B). However, a higher number of *Pht* genes was observed in Weining compared to Lo7 on chromosomes 4R, 6R and 7R, while on chromosome 5R only one *Pht* gene was discovered in Weining, compared to two in Lo7. These findings reveal an unequal distribution of phosphate transporter genes within the *Secale cereale* L genome.

**Figure 1.**
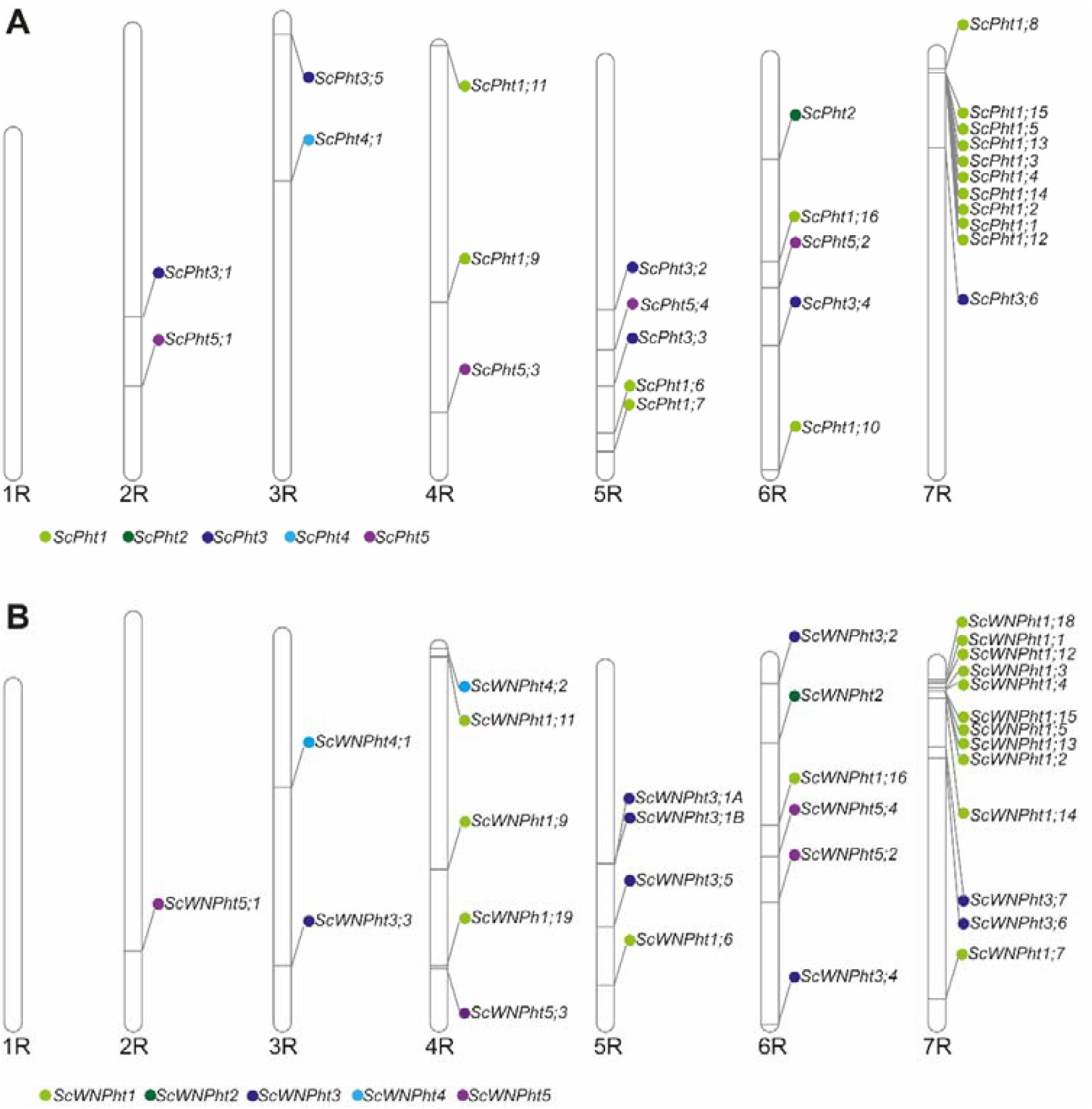
Distribution of *ScPht* genes on the rye Lo7 and Weining chromosomes. A) Overview of the seven rye Lo7 chromosomes and the *ScPht* genes location. B) Overview of the seven Weining Lo7 chromosomes and the *ScPht* genes location.

### Phylogenetic analysis of the family of PHT transporters

We constructed an unrooted phylogenetic protein tree using the neighbor-joining (NJ) method and containing a total of 139 PHT sequences from Arabidopsis (22 members), rice (27), sorghum (26) and both rye Lo7 (29), and Weining (35) to investigate the evolutionary relationships among PHT transporters. In our phylogenetic analysis, we included protein sequences from plants with reports of at least four PHT transporter families. We did not include OsPHT4;2 (LOC_ Os05g37820), OsPHT4;5 (LOC_ Os09g38410), and OsPHT4;6_2(LOC_ Os12g07970) since we could not retrieve these sequences from the current rice genome MSU release 7. Our results showed that the members of the same PHT family from different plants cluster together, forming five clades (Figure 2). The largest clade - PHT1 - contains 98 members and a rye-specific subgroup contains 16 PHT1 members. The PHT2 clade is the smallest and comprises four members.

**Figure 2.**
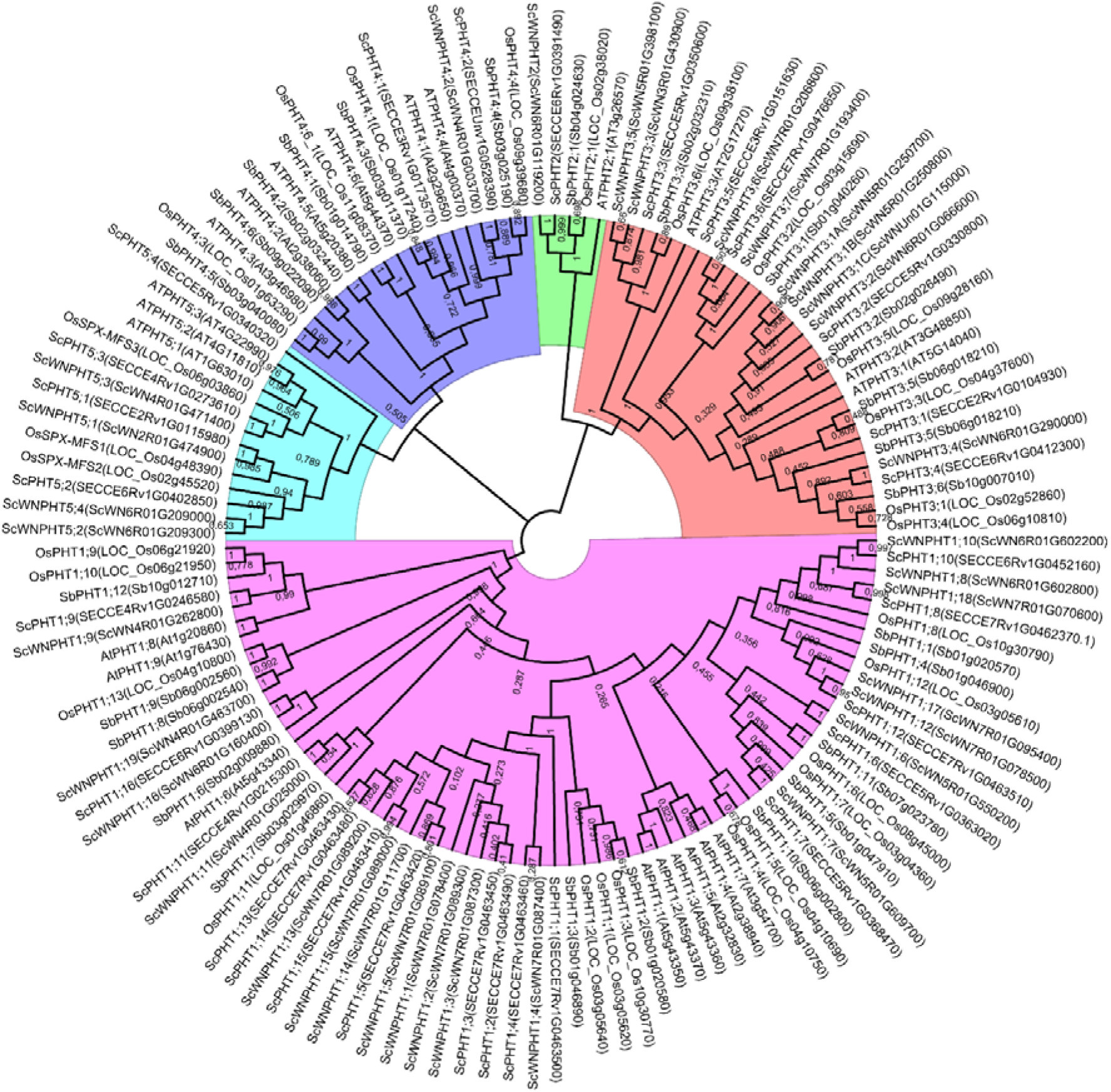
Phylogenetic relationships of PHT protein families from *Secale cereale* L. Lo7 and Weining, *Oryza sativa, Sorghum bicolor*, and *Arabidopsis thaliana*. The protein tree was constructed using the Neighbor-Joining method. The bootstrap consensus tree was inferred from 1000 replicates.

We further evaluated the PHT1 phylogenetic relationships among grasses to predict the potential function of rye PHT1 transporters (Figure 3). Our analysis showed that PHT1 transporters form 12 clusters. Most Arabidopsis PHT1 transporters – six out of the nine AtPHT1 – group into cluster I. Cluster II does not hold any *Triticeae* PHT1, while clusters VI and XII do not include rye phosphate transporters. Our analysis indicated that cluster III holds the most ScPHT transporters and contains almost exclusively *Triticeae* PHT1, except for brachypodium PHT1.3.

**Figure 3.**
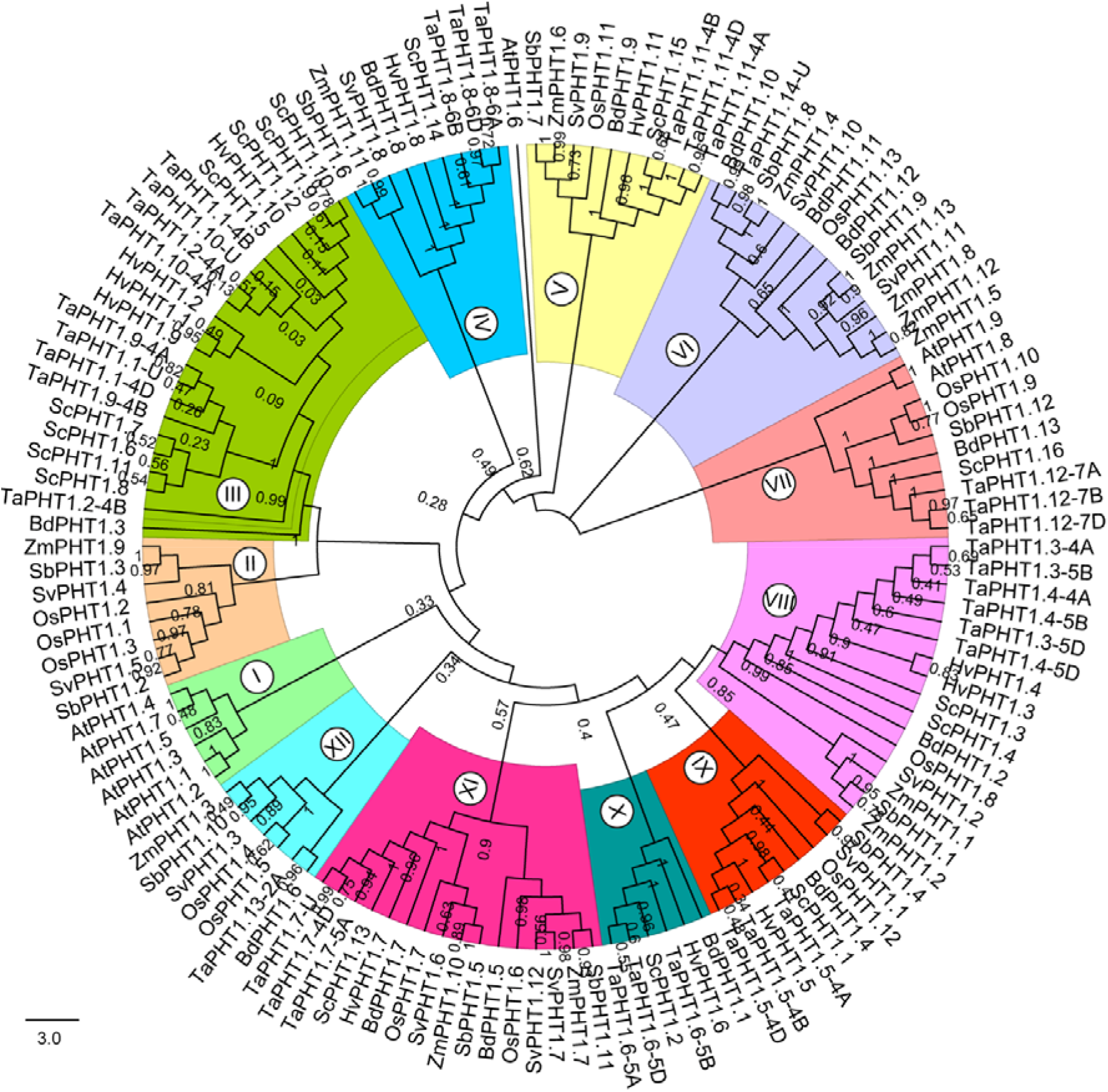
Phylogenetic relationships of PHT1 protein from different monocot grass species. The phylogenetic protein tree contains PHT1 protein from *Secale cereale* L., *Triticum aestivum, Hordeum vulgare, Oryza sativa, Sorghum bicolor, Zea mays, Setaria viridis, Brachypodium distachyon*, and Arabidopsis thaliana. The protein tree was constructed using the Neighbor-Joining method. The bootstrap consensus tree was inferred from 1000 replicates.

### Identification of the *cis*-acting regulatory elements

We analyzed the 2 kb upstream region of every putative rye *Pht* gene to gain insight into the potential transcriptional regulation during P starvation. We identified at least one PHR1-binding sequence (P1BS) in the promoter region of 13 *ScPht1* (Figure 4). *ScPht1;5, ScPht1;7*, and *ScPht1;15* promoters contain the highest number of P1BS *cis*-elements. The only *ScPht2* family member does not hold any P1BS *cis*-element. In the *ScPht3* family, *ScPht3;4, ScPht3;1*, and *ScPht3;5* contain two, one, and one P1BS *cis*-element, respectively. As for *ScPht*4 and *ScPht5* families, we found the phosphate deficiency responsive regulatory element in only one member (*ScPht4;2* and *ScPht5;3*) of each family. The presence of P1BS *cis*-elements suggests that P starvation regulates the expression of some members of each *ScPht* gene family.

**Figure 4.**
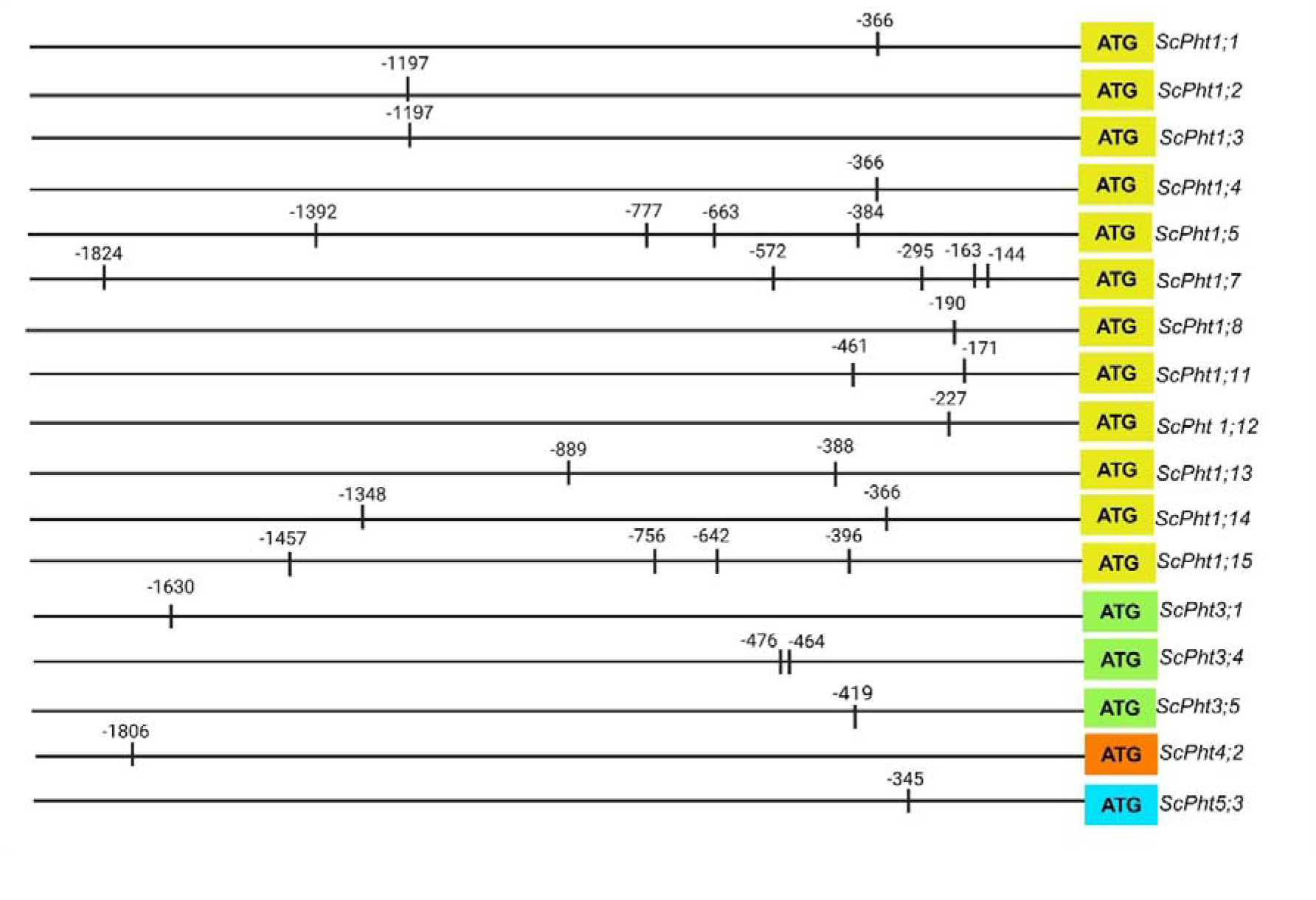
Location of P1BS *cis*-regulatory elements in the promoter region of *ScPht*s genes.

### Expression profiles of rye phosphate transporters in leaves and roots

Quantitative RT-PCR was used to analyze the responses of *ScPht* genes to Pi deficiency at two time points 14 and 21 days in the inbred line Lo7. We selected *ScPht1;6, ScPht1;7*, and *ScPht1;11* for primer design because they contain P1BS cis-elements and exhibit lower sequence similarity to each other within the *ScPht1* family: *ScPht1;6* shares 77,8% similarity with *ScPht1;7* and 69,13% with *ScPht1;11* (Additional file 1: Table S5). *ScPht3;4* and *ScPht3;5* have a higher sequence similarity of 84.69 %. *ScPht3;1* shares 75.25 % similarity with *ScPht3;4* and 72.30% similarity with *ScPht3;5*. Additionally, *ScPht5*.*1* and *ScPht5*.*3* have 40,61% sequence similarity, which allowed for the design of more specific primers. Primers specific to the following genes: *ScPht1;6, ScPht1;7, ScPht1;11, ScPht2, ScPht3;1, ScPht3;4, ScPht3;5* and *ScPht5;3* were designed (Additional file 1: Table S6). Only *ScPht1;6*; *ScPht*2, and *ScPht3;1* primers demonstrated high efficiency.

The relative expression of these three genes was examined across leaf and root tissues. In leaf tissue *ScPht1;6*; *ScPht2*, and *ScPht3;1* demonstrated different expression patterns (Figure 5A). Specifically, *ScPht1;6* showed statistically significantly higher expression under Pi deficiency at 14 and 21 days, while *ScPht2* exhibited significantly lower expression levels under Pi deficiency compared to control conditions at both 14 days and 21 days. For *ScPht3;1*, low expression levels were observed at both time points and the differences between treatments were not statistically significant.

**Figure 5.**
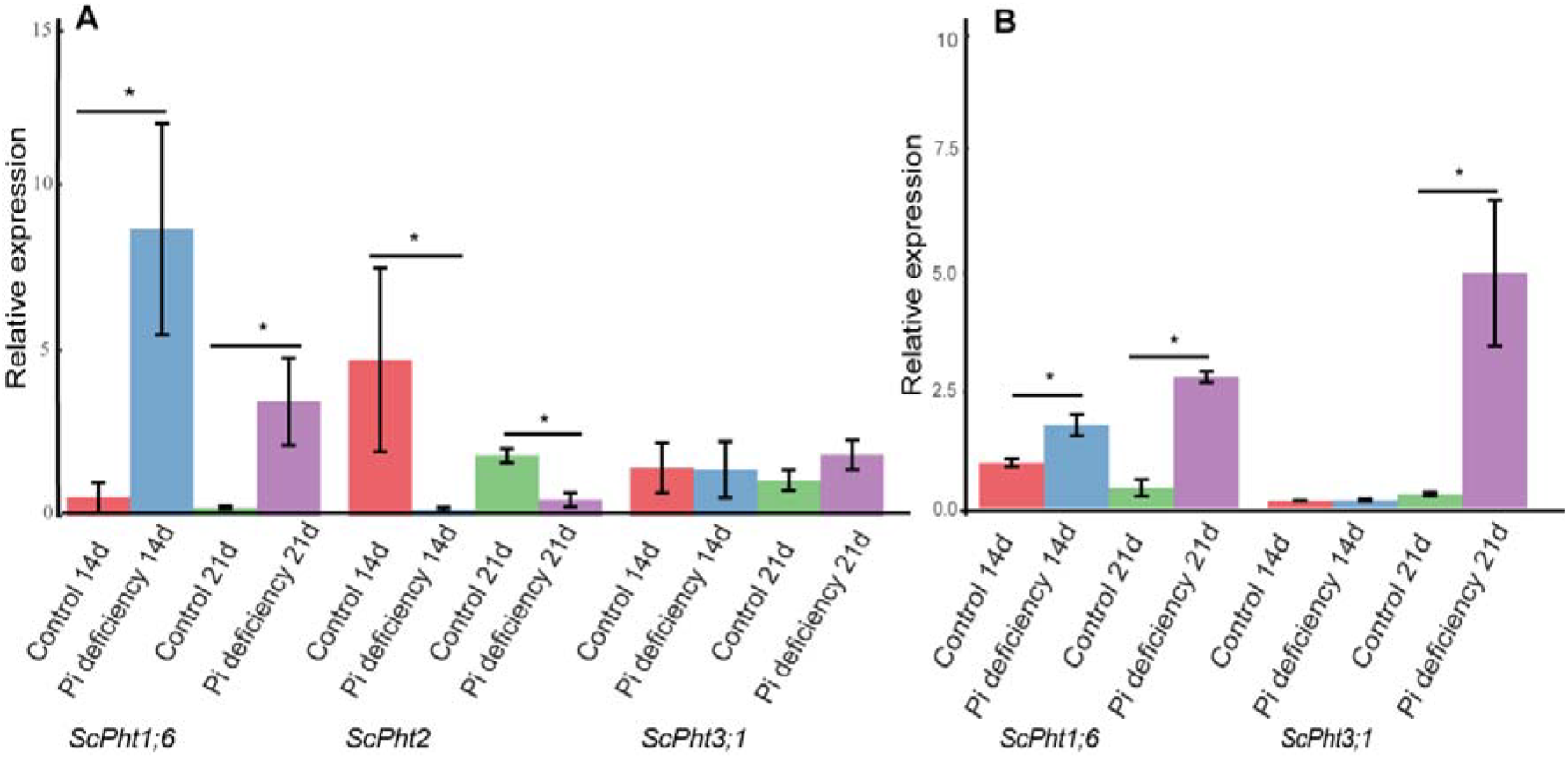
Relative expression levels of phosphate transporter genes assessed by qRT-PCR in leaf (A) and root (B) tissue under two treatment conditions: phosphate deficiency (Pi deficiency) and phosphorus sufficiency (control), at two-time points (14 days and 21 days). Each bar plot represents the mean 2^-ΔΔCt values obtained from three independent biological replicates, with error bars indicating the standard error of the mean (SEM). Statistical significance between control and treatment conditions was determined using the Kruskall test (*p < 0.05).

Only *ScPht1;6* and *ScPht3;1* genes exhibited differential expression patterns in root tissues (Figure 5B). *ScPht*2 transcripts were not detected in root tissue. Specifically, the expression level of *ScPht1;6* was significantly upregulated under Pi deficiency at 14 and 21 days. In the case of *ScPht3,1* a statistically significant upregulation of expression was observed in Pi deficiency samples at 21 days timepoint. The expression levels of *ScPht3;1* at 14 days were very low in both control and Pi deficiency samples and statistically significant differences were not observed.

### Low coverage resequencing (DArTreseq): phylogenetic relationships between 94 diverse rye accessions and sequence diversity of *ScPht* transporters

Initially, ca. 2,5 Mio. SNPs differentiating the 94 rye accessions included in the study were identified as a result of DArTreseq genotyping. After quality filtering 190 430 SNPs remained and were used for PCoA and NJ analyses. A high diversity of the collection was revealed (Figure 6A and B). Rye inbred lines separated from the remaining accessions, while landraces overlapped partially with cultivars, but displayed much higher diversity.

**Figure 6.**
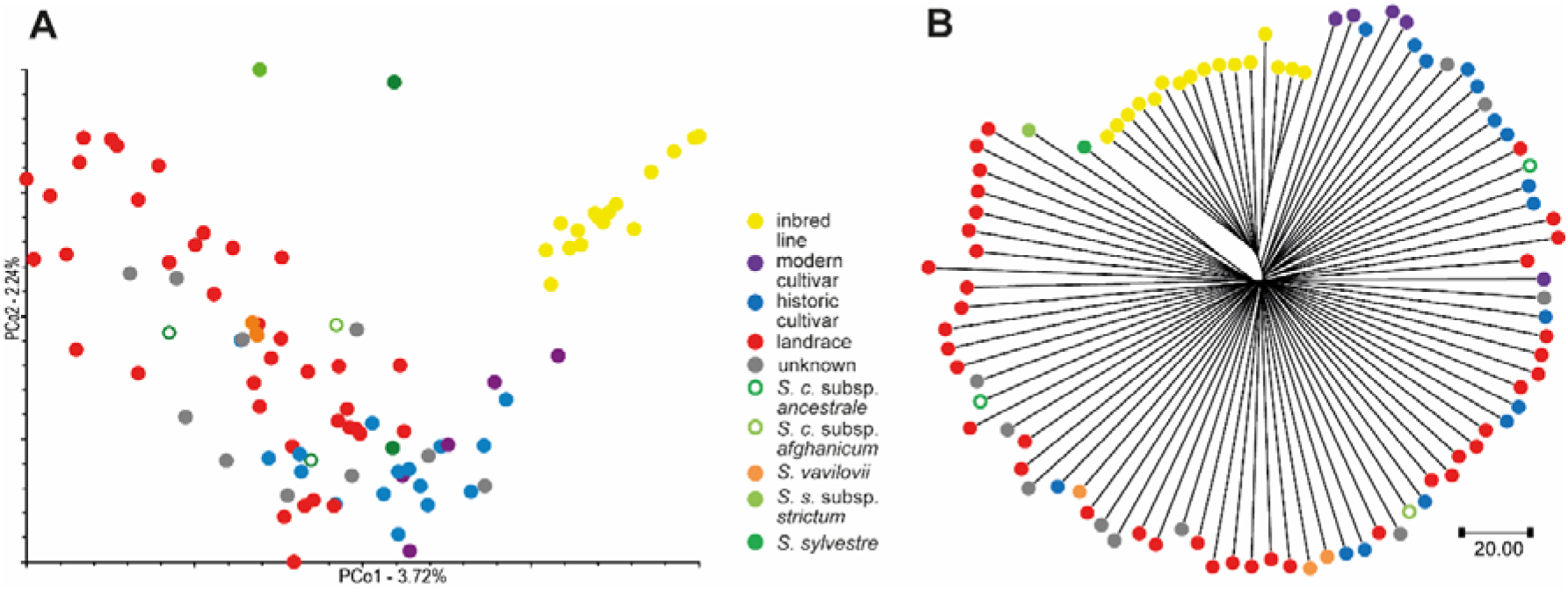

In total 820 polymorphic sites were observed within the putative 29 *ScPht* genes identified in the Lo7 genome. All identified polymorphisms were biallelic SNPs, with exception of two loci, where two alternative alleles were discovered. Transitions (G to A, C to T, A to G and T to C changes) constituted almost 54% of the discovered polymorphisms. The cumulative length of the analyzed genes and their putative regulatory regions was 185 414 bp, which corresponds to overall SNP density 1 SNP every 226 bp. The highest SNP density was observed in introns (one SNP per 128 bp), followed by CDS (one SNP per 187 bp) and UTRs (one SNP per 297 bp on average) (Additional File 1: Table S7). No polymorphisms were identified in the genes *ScPht1;2, ScPht1;3, ScPht1;4* and *ScPht1;14*. Within the remaining genes the SNP density ranged from one SNP per 84 bp for *ScPht1;6* to one SNP per 1341 bp of total gene length for *ScPht2*. The overall highest density of SNPs was observed in introns of *ScPht3;1* (one SNP per 84 bp) and the lowest – in UTRs of *ScPht4;1* (one SNP per 4000bp). Within the putative CDSs no SNP were identified in eight *ScPht1* family members – *ScPht1;1* - *ScPht1;5* and *ScPht1;13 - ScPht1;15*. Within the CDS of the remaining genes the SNP density ranged from one SNP per 60 bp for *ScPht5;4* to one SNP per 848 for *ScPht2*.

The number of polymorphic sites within *ScPht* genes per accession ranged from 51 (in inbred line L318) to 201 in the landrace NSL308 from Turkey, 131 on average. The number of accessions with a given variant ranged from 1 to 70, on average a variant occurred in 15 accessions. In total 11 private variants (occurring only in a single accession from the set) were identified. Six of the private variants occurred in wild/weedy accessions, four in landraces, and one in a historic cultivar. The accessions OBW059 (wild/weedy - S. s. subsp. *strictum*) and NSL282 (a landrace from Pakistan) contained two private variants each, while each of the remaining seven private variants occurred in a different accession. The private variants were identified in ten genes: four *ScPht1* genes (*ScPht1;1, ScPht1;5, ScPht1;9* and *ScPht1;11*), three *ScPht3* genes (*ScPht3;1, ScPht3;2* and *ScPht3;4*), one *ScPht4* gene (*ScPht4;1*), and two *ScPht5* genes (*ScPht5;1, ScPht5;2*). A single private variant was found in each of these genes, with exception of the *ScPht1;11*, where two private variants were discovered. The majority of private variants were located in UTR regions, only the private variants in *ScPht3;1* and *ScPht4;1* were located in introns. In total 12 putatively deleterious variants were identified by SIFT4G analysis in seven *ScPht* genes including nine deleterious variants with a low confidence score (Additional File 1: Table S8). Overall the deleterious variants were identified mainly in the *ScPht1* family members: two variants in each of the genes *ScPht1;7, ScPht 1:12, ScPht 1;16* and one variant in each of the genes *ScPht1;8, ScPht1;9, ScPht1;10, ScPht3;2, ScPht5;3* and *ScPht5;4*. In the analyzed set the number of accessions with a given deleterious variant ranged from three to 26. The location of high confidence deleterious SNPs and corresponding amino acid changes is shown in Figure 7.

**Figure 7.**
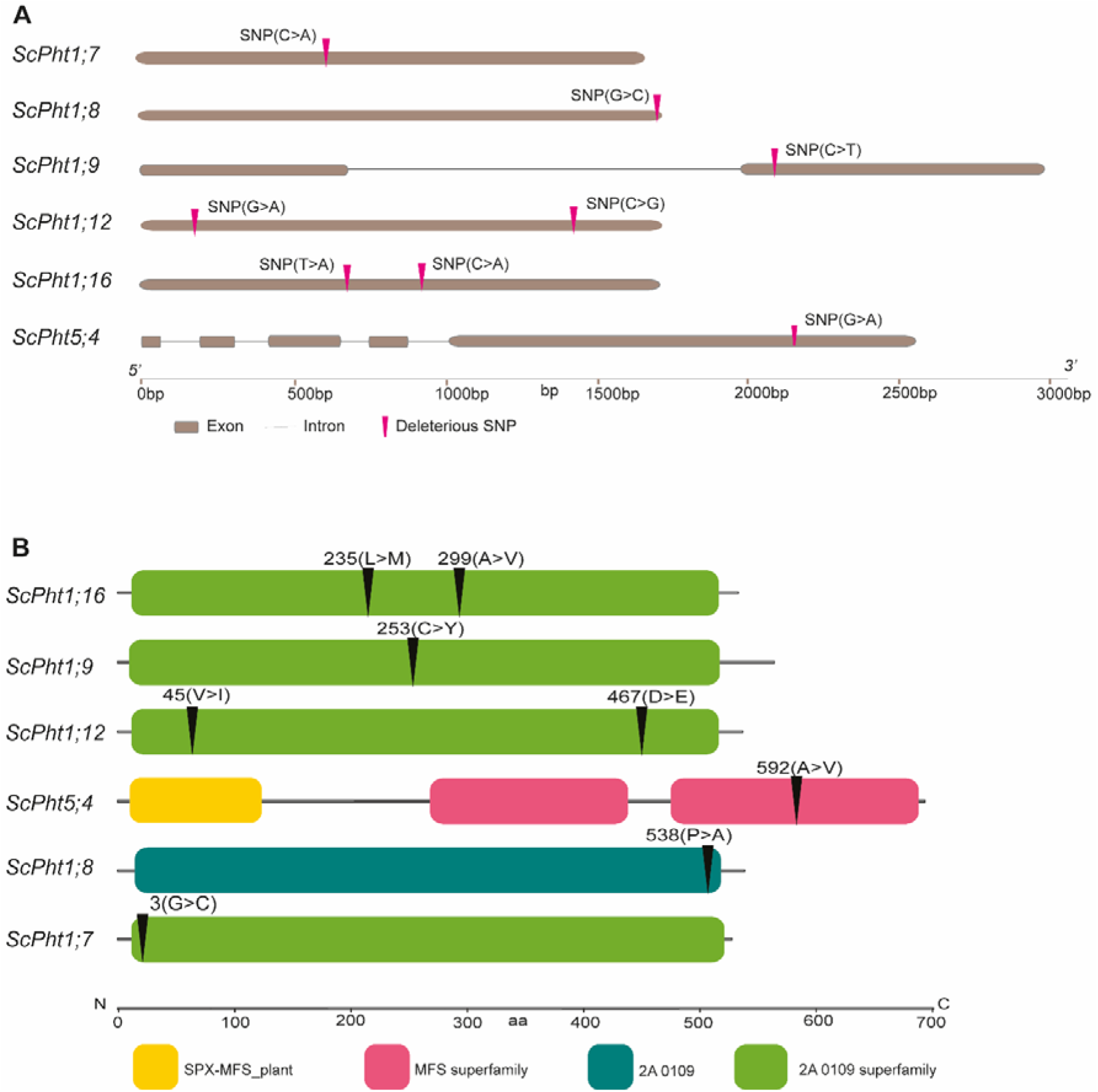
Genetic diversity of 94 rye accessions analyzed by DArTreseq. A) PCoA plot based on 190430 genome-wide SNPs. B) NJ tree based on 190430 genome-wide SNPs.

**Figure 8.** Diagram of the deleterious SNP location within the *ScPht* members. A) Location of deleterious SNP within the coding regions of *ScPht* genes. B) Amino acid changes due to deleterious SNPs in the protein sequences of PHT transporters.

## DISCUSSION

The distribution of phosphate to the various plant tissues and cellular compartments depends on PHT transporters. In our study, we surveyed rye Lo7 and Weining reference genomes to identify members of the *Pht* transporter families. The number of *Pht* genes in each rye reference genome – 29 and 35 in Lo7 and Weining, respectively – differs, particularly among the *Pht1* and *Pht3* family members. These discrepancies in the *Pht* gene numbers between reference genomes could be due to genomic diversity or *de novo* genome sequencing approaches.

Genetic diversity within and between species often arises from small (e.g. single nucleotide polymorphism, insertion/deletions) and large-scale polymorphism [57]. The large-scale polymorphisms, also known as structural variations (SVs), come in many forms – copy number variation (CNV), insertions, deletions, duplications, inversions, intra and interchromosomal translocation – and sizes – larger than 50 bp to several Mb [58].

Several comparative genomic studies in crop species (e.g., rice, wheat, barley, maize) have uncovered structural variation – specifically the presence or absence of loci – conferring advantageous agronomical and adaptive traits [59, 60]. For instance, *SNORKEL1* and *SNORKEL2* genes are integral parts of the genetic makeup of the deepwater rice cultivar, but both loci are absent in cultivars unfit to grow in flooding conditions [61]. Similarly, the *Pup1* locus – present in the aus-type Kasalath rice variety but missing in the Nipponbare cultivar – provides advantages under low-P environments by influencing the root architecture [36, 62]. These examples point out the risk of overlooking genetic variants associated with adaptive traits when relaying in a single reference genome.

In rye, large inversions occur in various chromosomes among members of the *Secale* genus. Three-dimensional conformation capture sequencing (Hi-C) revealed large inversion in chromosomes 1R, 2R, 4R, and 5R between rye Lo7 and Lo255 inbred lines [37]. Large inversions are evident in every chromosome when comparing Lo7 with *Secale strictum*, a distant relative of *Secale cereale*. The phylogenetic distance between the rye inbred line Lo7 and other members of the *Secale* genus dictates the number of large inversions among species [37]. However, precise data on the extent of structural variation regarding the presence/absence variation of gene loci are not available yet for rye. The variation in the *Pht* gene number between rye Lo7 and Weining reference genomes might reflect the adaptation to their local environments.

The Lo7 and Weining genomes were sequenced and assembled following different strategies. Both rye genomes were assembled to the chromosome level using Hi-C and BioNano optical mapping [37, 41]. The genome assembly strategies between rye Lo7 and Weining contrasted on the choice of sequencing platforms: Illumina short reads and PacBio for rye Lo7 and Weining, respectively [37, 41]. Although Illumina sequencing platforms have been the workhorse of several de novo genome sequencing projects, the short-length read imposes challenges when assembling large and complex plant genomes [63]. A large content of repetitive sequences frequently characterizes large plant genomes, which in rye represent around 90% of the genome [64, 65]. Moreover, large gene families in plant genomes add a layer of difficulty to genome assembly [65]. As shown in our study, many members of the *Pht1* family – in both Lo7 and Weining genomes – share a high nucleotide sequence similarity (> 90 %) among them. Thus, the short-sequencing-based Lo7 genome nature might have underestimated the numbers of *Pht1* family members. A similar situation might be the cause of the *Pht3* family members’ variation between Lo7 and Weining genomes. For instance, the number of *Pht3* members in the Weining chromosomes 5R, 6R, and 7R doubles the number of *Pht3* genes in the same chromosomes of the rye Lo7. Consequently, we cannot rule out the inherent characteristics of the sequencing platform and assembly approaches as a source of *Pht* gene number variations.

We also observed variation in the number of members from each *Pht* family across plant species. For instance, both rye reference genomes – 16 *Pht1* in Lo7 and 19 *Pht1* in Weining – hold a larger number of *Pht1* genes compared to Arabidopsis (9 *Pht1*), rice (13 *Pht1*), barley (11), maize (13), brachypodium (13), *Setaria* (12), and sorghum (12 *Pht1*).

The expansion of gene families is often attributed to fragment and tandem duplication as well as transposable replication and whole genome duplication (WGD) events [66]. The rye Weining genome contains a higher number of proximal and tandem duplication genes compared to other grasses - barley, diploid wheat, *Aegilops tauschii*, brachypodium, and rice [41]. Similarly, the transposed duplicated gene number in rye Weining is considerably larger than in diploid wheat and *Aegilops tauschii*. Some rye Weining genes involved in starch biosynthesis had undergone transposed (*ScSSIV, ScDPEI, ScSuSy1, ScSuSy1*, and *ScUDPaseI*), tandem (*ScPHO2*), proximal (*ScAGP-L2-p, and ScSBE1*), dispersal (*SSIIIa*) duplication events [41]. In the rye Lo7 genome, the number of members of mildew-resistant (*Pm2, Pm3*, and *Mla*) gene families varies compared to other members of the *Triticeae* tribe [37]. For instance, the rye Lo7 *Mla* family contains ten members while the barley MLA family has four.

Through our phylogenetic analysis (Figure 3), we could hypothesize about the role of rye PHT in P homeostasis, since several PHT1 transporters have been experimentally validated. For instance, cluster III contains various wheat (TaPHT1.1, TaPHT1.2, TaPHT1.9, and TaPHT1.10) and barley (HvPHT1.1) transporters exhibiting Pi-transport activity, and increasing expression in roots under Pi deficiency [19, 67]. The PHT1 phylogenetic protein tree also revealed interesting relationships between rye, maize, and rice phosphate transporters. For example, the rye ScPHT1;6 closely relates to HvPHT1.6. The barley PHT1;6 and rice PHT1;7 transporters have been suggested to remobilize stored Pi from old to young leaves [67, 68]. Moreover, the putative rye ScPHT1;9 transporter groups with the functionally redundant rice OsPHT1;9 and OsPHT1;10 – both transporters are involved in Pi-uptake [69]. The cluster V groups ScPHT1;11 with OsPHT1;11 and ZmPHT1;6 – both transporters are known to be involved in mycorrhiza-dependent Pi-uptake [26, 70].

To date, there is no report about the expression profiles of the genes involved in response to P deficiency in rye. We aimed to address this knowledge gap by evaluating the gene expression of *ScPht* family members in leaves and roots under Pi-deficient conditions. We initially selected *Pht* genes for qRT-PCR analysis based on the presence of P1BS *cis*-elements and whether we could design gene-specific primers. The high sequence similarity among *Pht* genes has limited our ability to generate primers to measure the expression of the individual genes under Pi deficiency. Noteworthy, we included *ScPht2* in our analysis despite it lacking a P1BS *cis*-element. This inclusion is based on functional evidence concerning its orthologs rather than the presence of a specific regulatory element, highlighting the gene’s critical role in Pi transport mechanisms across different species - Arabidopsis, wheat, and sorghum [71–73].

Under Pi-starvation stress, the expression of *Pht1* genes was shown to be significantly upregulated to enhance the roots’ ability to absorb Pi from the soil and to facilitate Pi remobilization within the plant [74, 75]. Additionally, the transcript levels of *Pht1* genes have been correlated with P utilization efficiency in barley [76]. In this context, our study examined the expression of *ScPht1;6*. The results revealed a statistically significant expression upregulation in root samples subjected to Pi deficiency compared to the control samples at 14 and 21 days (Figure 5B). *ScPht1;6* also exhibited a statistically significant expression upregulation in Pi-deficient leaves samples compared to control samples at both 14 and 21 days, with the highest expression levels observed at 14 days of Pi deficiency. These results indicate that while the gene remains expressed, its transcript levels decrease as the duration of Pi deficiency increases. Our phylogenetic analysis identified *ScPht1;6* as an ortholog of *TaPht1;6-* 5A in wheat (Figure 3). Ta*Pht*1;6-5A exhibited differential organ-specific expression, enhancing Pi acquisition and accumulation in all organs of a Pi-efficient wheat cultivar under Pi withdrawal [77]. Notably, *TaPht1;6* was the only *TaPht1* transporter with high transcript abundance in the ear and rachis during Pi starvation [78]. Similarly, *OsPht1;6*, the rice ortholog of *ScPht*1;6 (Figure 3), exhibited significant upregulation after 14 days of low P conditions [79]. In barley, *HvPht1;6*, an identified ortholog of *ScPht1;6* (Figure 3) showed significantly elevated transcript levels at 17 days under severe Pi deficiency in shoots and roots [76]. This similarity in upregulation patterns under Pi deficiency supports the hypothesis that *Pht1* transporters are crucial for adaptation to low Pi availability [80]. Our findings with *ScPht1;6* align with these observations in both tissues, where significant expression upregulation was noted under Pi deficiency.

In a similar vein, we examined the expression of *ScPht2*, the sole identified gene of this family. We found that *ScPht2* was not expressed in roots but displayed significant downregulation in leaves under Pi deficiency. Thus, *ScPht2* demonstrated tissue-specific expression in leaves, comparable to Arabidopsis and rice, which is considered to imply their conserved function in Pi transport into chloroplasts in those plants [71, 81]. In our experiments, *ScPht2* expression remains low with no significant variations over time in P-deficient conditions. These results suggest that *ScPht2* may not play a direct role in Pi uptake during Pi stress but could be essential for maintaining Pi homeostasis under normal conditions. [77] found that the expression profile of *TaPht2;1* varies with genotype and is linked to plant P efficiency in wheat. They proposed that *TaPht2;1* could serve as a marker gene for identifying genotypes with high P efficiency. Another study in wheat revealed that the expression level of *TaPht2;1* was significantly enhanced in leaves when subjected to Pi deficiency [72]. In rice, the expression of *OsPht2;1* was induced in leaves but not in roots under Pi deficiency [81]. Similarly to *ScPht2, AtPht2;1* and *TaPht2;1* were mainly expressed in the shoots. Additionally, the *CsPht2* genes were preferentially expressed in leaves during the early and vegetative stages in response to Pi deficiency in *Camelina sativa* [82], Likewise, the expression of two *GmPht2* genes was induced by low-Pi stress in soybean leaves [83]. In a similar manner to our results, *CaPht2;1* showed minimal expression in shoots during Pi stress in *Capsicum annuum* [84]. Overall, these findings underscore the diverse yet tissue-specific transcriptional regulation of *Pht2* genes in Pi deficiency across different plant species.

The plant gene family *Pht3* is known to localize in the mitochondrial inner membrane [85]. In our study, *ScPht3;1* expression levels in leaves showed no significant differences between treatments at both time points, indicating it is not dependent on Pi availability. Conversely, *ScPht3;1* expression levels in roots displayed a significant upregulation at 21 days of Pi deficiency, indicating a responsive adaptation to prolonged Pi deprivation in the root system. In sorghum, *SbPht3;6*, which is an ortholog of *ScPht3;1*, was upregulated more than 2-fold in the leaves and downregulated in roots after 14 days of Pi starvation [73]. In pepper, a differential expression of *CaPht3*, a putative ortholog of *ScPht3;1*, was observed in both roots and leaves during P deficiency [84]. In contrast, our findings regarding *ScPht3;1* expression levels suggest a specific function in Pi -deficiency response in roots.

Our qRT-PCR data indicated that the analyzed *ScPht* genes are Pi-deficiency responsive (Figure 5). To better characterize functions of the identified *ScPht* genes in Pi uptake and translocation, further research is needed, involving examination of the Pi affinity of ScPHT proteins and gene functional analyses.

We have analyzed the genetic diversity of the identified *ScPht* genes in a collection of 94 rye accessions with various improvement status and geographic origins using low coverage resequencing (DArTreseq). Accessions originating from different cultivation environments were included in the set, for example, landraces /historical cultivars from areas such as Brazil, Norway, Finland or Sweden, where the very high P-retention potential of the soil results in a low availability of P for plants [86], as well as modern varieties bred for cultivation in Central Europe (Germany, Poland), where P supply is not a limiting factor.

Neighbour Joining and Principal Coordinates analyses revealed a large diversity of the collection. The obtained picture of genetic diversity patterns is in good agreement with results of previous studies on rye genetic diversity [43, 87–89] which indicated genetic distinctness of rye inbred lines, broad diversity of genetic resources and narrower diversity of historical and modern rye varieties. Therefore it can be assumed that the collection is representative and covers a broad spectrum of rye genetic diversity. Nevertheless, the average SNP density observed in this study across the *ScPht* genes (1 SNP every 226 bp) was much lower than reported earlier for rye by Hawliczek et al. [43] - one SNP or InDel every 12 bp. This striking difference can result from methodological differences between studies. Since rye has a very large (8Gb) genome and a high level of heterozygosity and heterogeneity is typical for rye accessions, in the present study, for each accession a single plant was analyzed, to facilitate the SNP calling. Hence, the within accession diversity was not probed. In the study by Hawliczek et al. [43], sample pooling was deployed and each accession was represented by 96 plants, while the use of amplicon sequencing approach ensured a very high sequencing coverage. Therefore a very precise assessment of intra-accession diversity was achieved. Those are likely the key factors that contributed to the much higher SNP density discovered by Hawliczek et al. [43]. Nevertheless, the average SNP density observed in this study is also several fold lower than in other earlier studies on rye SNP diversity, where intra-accession diversity was not taken into regard: 1 SNP/52 bp [90], 1 SNP/58 bp [91]. Although the technical limitation of the applied approach of probing the gene diversity (low coverage resequencing in a large genome species) could have contributed to the lower SNP density to some extent [92], the observed differences in SNP density could also reflect differences in mutation rates across the genomic regions analyzed in the studies mentioned above. It has been observed that mutation rates vary in plant genomes, with differences occurring across genotypes, genome locations, gene functions, etc. [93] and it is assumed that the coding sequences of developmental genes are strongly conserved, whereas genes with roles in defense evolve more rapidly [94]. While *phosphate transporter* genes are the sole focus of the present study, the earlier analyses involved: six genes related mostly to biotic stress resistance and seed quality [43], 12 genes with putative roles in frost tolerance [90], 14 ESTs with various putative functions [91].

Furthermore, SNP density varied markedly within each *ScPht* family: for example, the largest, *ScPht1* family comprised genes with SNP density ranging from 1 SNP/84 bp to 1 SNP/795 bp, as well as three genes where no polymorphism was detected in the 94 rye accession studied. These findings might reflect the evolutionary history of these gene families, involving recurrent gene duplication and accumulation of genetic diversity in some copies by positive natural selection or other mechanisms [95, 96]. Moreover, different mutation rates can be a consequence of such factors as chromosomal location, gene orientation or transcriptional activity [97].

In total, we identified 820 polymorphic sites within *ScPht* genes based on low-coverage resequencing *via* DArTreseq of 94 rye accessions, among them 11 were private variants. The vast majority of those private variants occurred in wild accessions or landraces, which further confirmed the value of crop wild relatives and landraces as a source of potentially useful variation in rye and other crops [43, 98, 99]. It was shown in the past that landraces, which constitute ca. 41 %of our germplasm set, are an important source of genes enhancing nutrient uptake and utilization, as well as adaptation to water, salinity and high temperature stresses [98]. Furthermore, 12 of the variants were found to be deleterious to gene function. Therefore, it can be argued that, despite certain shortcomings (low coverage, disregarding intra-accession diversity), the adopted approach of detecting sequence diversity (DArTreseq) allows for a quick and relatively cost-effective insight into the diversity of all annotated genes, and produces data which permit for relative comparison of polymorphism levels across genes/gene families of interest and quick identification and prioritization of candidates for further functional studies.

## Conclusions

Rye genome contains putative phosphate transporter genes from all known *Pht* families. Phylogenetically, putative rye transporters exhibit a close relationship to previously functionally characterized PHT proteins from other grasses. The two available rye reference genomes differ in the number of *Pht*1 and *Pht*3 genes and the polymorphism level varies markedly among the identified *ScPht* genes within the rye diversity panel, suggesting multiple layers of adaptation to local environments existing in rye. *ScPht1;6, ScPht2* and *ScPht3;1* genes are responsive to Pi-deficiency, which might suggest their involvement in Pi homeostasis. DArTreseq genotyping permits for a quick and cost-effective assessment of polymorphism levels across genes/gene families and supports identification and prioritization of candidates for further studies.

Our report is the first step toward elucidating the mechanisms of rye’s response to Pi deficiency. Collectively our findings provide the foundation for selecting most promising candidates for further functional characterization.

## DECLARATIONS

### Ethics approval and consent to participate

All methods were carried out in accordance with relevant guidelines and regulations.

### Consent for publication

Not applicable

### Availability of data and materials

FASTQ files related to DArTreseq analysis of 94 rye accessions generated and analyzed in this study are available in NCBI BioProject PRJNA1091674.

## Competing interests

The authors declare that they have no competing interests.

## Funding

This research was funded by the National Science Centre, Poland grant No. 2020/37/B/NZ9/00738

## Authors’ contributions

DCR, BWK, SW, FA and HAA performed the experiments, DCR, BWK, SW, BJT, AH, MK and JM analyzed the data, HBB and BJT contributed data/analysis tools, DCR and HBB conceived, designed and supervised the experiments, DCR, HBB, SW and BWK wrote the manuscript, BJT reviewed and edited the manuscript. HBB acquired funding. All authors read and approved the final manuscript.

## Acknowledgments

Authors would like to thank Prof. Beata Myśków from Department of Plant Genetics, Breeding and Biotechnology, West Pomeranian University of Technology (Szczecin, Poland) for providing seeds of rye inbred lines for this study.

## Additional Files

### Additional File 1

**Table S1**. Description of rye accessions analyzed using DArTreseq: name, source, geographic origin, and improvement status

**Table S2**. List of PHT transporters used to build up protein phylogenetic trees.

**Table S3**. List of putative rye *Pht* genes identified in the Lo7 genome and the features of each PHT transporter.

**Table S4**. List of putative rye *Pht* genes identified in the Weining genome and the features of each PHT transporter.

**Table S5**. Percentage of sequence similarity between *ScPht* gene families’ members.

**Table S6**. Primer sequences used for real-time qPCR analysis of *ScPht* transporters expression.

**Table S7**. SNP densities in *ScPht* genes based on low coverage resequencing (DArTreseq) of 94 rye accessions.

**Table S8**. Deleterious variants in *ScPht* genes predicted by SIFT4G.

## Notes

### Competing Interest Statement

The authors have declared no competing interest.

## REFERENCES

1. Chithrameenal K, Alagarasan G, Raveendran M, Robin S, Meena S, Ramanathan A, et al. Genetic enhancement of phosphorus starvation tolerance through marker assisted introgression of OsPSTOL1 gene in rice genotypes harbouring bacterial blight and blast resistance. PLoS One. 2018;13:e0204144.

2. Ringeval B, Augusto L, Monod H, van Apeldoorn D, Bouwman L, Yang X, et al. Phosphorus in agricultural soils: drivers of its distribution at the global scale. Glob Chang Biol. 2017;23:3418–32.

3. Hou E, Luo Y, Kuang Y, Chen C, Lu X, Jiang L, et al. Global meta-analysis shows pervasive phosphorus limitation of aboveground plant production in natural terrestrial ecosystems. Nat Commun. 2020;11:1–9.

4. Penn CJ, Camberato JJ. A critical review on soil chemical processes that control how soil ph affects phosphorus availability to plants. Agric. 2019;9:1–18.

5. van de Wiel CCM, van der Linden CG, Scholten OE. Improving phosphorus use efficiency in agriculture: opportunities for breeding. Euphytica. 2016;207:1–22.

6. Mehra P, Pandey BK, Giri J. Genome-wide DNA polymorphisms in low Phosphate tolerant and sensitive rice genotypes. Sci Rep. 2015;5:13090.

7. Heuer S, Gaxiola R, Schilling R, Herrera-Estrella L, López-Arredondo D, Wissuwa M, et al. Improving phosphorus use efficiency: a complex trait with emerging opportunities. Plant J. 2017;90:868–85.

8. Młodzińska E, Zboińska M. Phosphate uptake and allocation - A closer look at arabidopsis thaliana L. And Oryza sativa L. Front Plant Sci. 2016;7:1–19.

9. Puga MI, Mateos I, Charukesi R, Wang Z, Franco-Zorrilla JM, De Lorenzo L, et al. SPX1 is a phosphate-dependent inhibitor of Phosphate Starvation Response 1 in Arabidopsis. Proc Natl Acad Sci U S A. 2014;111:14947–52.

10. Wang Z, Ruan W, Shi J, Zhang L, Xiang D, Yang C, et al. Rice SPX1 and SPX2 inhibit phosphate starvation responses through interacting with PHR2 in a phosphate-dependent manner. Proc Natl Acad Sci U S A. 2014;111:14953–8.

11. Liu TY, Huang TK, Yang SY, Hong YT, Huang SM, Wang FN, et al. Identification of plant vacuolar transporters mediating phosphate storage. Nat Commun. 2016;7.

12. Zhang C, Meng S, Li M, Zhao Z. Genomic identification and expression analysis of the phosphate transporter gene family in poplar. Front Plant Sci. 2016;7 September.

13. Wang D, Lv S, Jiang P, Li Y. Roles, Regulation, and Agricultural Application of Plant Phosphate Transporters. Front Plant Sci. 2017;8 May:1–14.

14. Wang Y, Wang F, Lu H, Liu Y, Mao C. Phosphate Uptake and Transport in Plants: An Elaborate Regulatory System. Plant Cell Physiol. 2021;62:564–72.

15. Nussaume L, Kanno S, Javot H, Marin E, Pochon N, Ayadi A, et al. Phosphate import in plants: Focus on the PHT1 transporters. Front Plant Sci. 2011;2 NOV:1–12.

16. Liu F, Chang XJ, Ye Y, Xie WB, Wu P, Lian XM. Comprehensive sequence and whole-life-cycle expression profile analysis of the phosphate transporter gene family in rice. Mol Plant. 2011;4:1105– 22.

17. Preuss CP, Huang CY, Gilliham M, Tyerman SD. Channel-like characteristics of the low-affinity barley phosphate transporter PHT1;6 when expressed in Xenopus oocytes. Plant Physiol. 2010;152:1431–41.

18. Liu F, Xu Y, Jiang H, Jiang C, Du Y, Gong C, et al. Systematic identification, evolution and expression analysis of the Zea mays PHT1 gene family reveals several new members involved in root colonization by arbuscular mycorrhizal fungi. International Journal of Molecular Sciences. 2016;17.

19. Teng W, Zhao Y-Y, Zhao X-Q, He X, Ma W-Y, Deng Y, et al. Genome-wide Identification, Characterization, and Expression Analysis of PHT1 Phosphate Transporters in Wheat. Front Plant Sci. 2017;8 April:1–14.

20. Ceasar SA, Hodge A, Baker A, Baldwin SA. Phosphate concentration and arbuscular mycorrhizal colonisation influence the growth, yield and expression of twelve PHT1 family phosphate transporters in foxtail millet (Setaria italica). PLoS One. 2014;9.

21. Walder F, Brulé D, Koegel S, Wiemken A, Boller T, Courty PE. Plant phosphorus acquisition in a common mycorrhizal network: Regulation of phosphate transporter genes of the Pht1 family in sorghum and flax. New Phytol. 2015;205:1632–45.

22. Maharajan T, Krishna TPA, Kiriyanthan RM, Ignacimuthu S, Ceasar SA. Improving abiotic stress tolerance in sorghum: focus on the nutrient transporters and marker-assisted breeding. Planta. 2021;254:1–16.

23. Pudake RN, Mehta CM, Mohanta TK, Sharma S, Varma A, Sharma AK. Expression of four phosphate transporter genes from Finger millet (Eleusine coracana L.) in response to mycorrhizal colonization and Pi stress. 3 Biotech. 2017;7.

24. Rausch C, Zimmermann P, Amrhein N, Bucher M. Expression analysis suggests novel roles for the plastidic phosphate transporter Pht2;1 in auto- and heterotrophic tissues in potato and Arabidopsis. Plant J. 2004;39:13–28.

25. Rausch C, Bucher M. Molecular mechanisms of phosphate transport in plants. Planta. 2002;216:23–37.

26. Zhu W, Miao Q, Sun D, Yang G, Wu C, Huang J, et al. The mitochondrial phosphate transporters modulate plant responses to salt stress via affecting ATP and gibberellin metabolism in Arabidopsis thaliana. PLoS One. 2012;7:1–10.

27. Guo B, Jin Y, Wussler C, Blancaflor EB, Motes CM, Versaw WK. Functional analysis of the Arabidopsis PHT4 family of intracellular phosphate transporters. New Phytol. 2008;177:889–98.

28. Liu Y, Wang L, Deng M, Li Z, Lu Y, Wang J, et al. Genome-wide association study of phosphorus-deficiency-tolerance traits in Aegilops tauschii. Theor Appl Genet. 2015;128:2203–12.

29. Bouain N, Doumas P, Rouached H. the Root System Response to Phosphate Deficiency in Arabidopsis. Curr Genomics. 2016;:308–14.

30. Ning L, Kan G, Du W, Guo S, Wang Q, Zhang G, et al. Association analysis for detecting significant single nucleotide polymorphisms for phosphorus-deficiency tolerance at the seedling stage in soybean [Glycine max (L) Merr.]. Breed Sci. 2016;66:191–203.

31. Lin Y, Chen G, Hu H, Yang X, Zhang Z, Jiang X, et al. Phenotypic and genetic variation in phosphorus-deficiency-tolerance traits in Chinese wheat landraces. BMC Plant Biol. 2020;20:1–9.

32. Luo B, Zhang G, Yu T, Zhang C, Yang G, Luo X, et al. Genome-wide association studies dissect low-phosphorus stress response genes underling field and seedling traits in maize. Theor Appl Genet. 2024;137:1–17.

33. Bouain N, Korte A, Satbhai SB, Nam HI, Rhee SY, Busch W, et al. Systems genomics approaches provide new insights into Arabidopsis thaliana root growth regulation under combinatorial mineral nutrient limitation. PLoS Genet. 2019;15.

34. Soumya PR, Burridge AJ, Singh N, Batra R, Pandey R, Kalia S, et al. Population structure and genome-wide association studies in bread wheat for phosphorus efficiency traits using 35 K Wheat Breeder’s Affymetrix array. Sci Rep. 2021;11:1–17.

35. Yan M, Feng F, Xu X, Fan P, Lou Q, Chen L, et al. Genome-wide association study identifies a gene conferring high physiological phosphorus use efficiency in rice. Front Plant Sci. 2023;14 March:1–12.

36. Gamuyao R, Chin JH, Pariasca-Tanaka J, Pesaresi P, Catausan S, Dalid C, et al. The protein kinase Pstol1 from traditional rice confers tolerance of phosphorus deficiency. Nature. 2012;488:535–9.

37. Rabanus-Wallace TM, Hackauf B, Mascher M, Lux T, Wicker T, Gundlach H, et al. Chromosome-scale genome assembly provides insights into rye biology, evolution and agronomic potential. Nat Genet. 2021;53:564–73.

38. Crespo-Herrera LA, Garkava-Gustavsson L, Åhman I. A systematic review of rye (Secale cereale L.) as a source of resistance to pathogens and pests in wheat (Triticum aestivum L.). Hereditas. 2017;154:14.

39. Geiger HH, Miedaner T. Rye breeding. In: Carena MJ, editor. Cereals (Handbook of Plant Breeding, volume 3). 1st edition. New York, NY: Springer US; 2009. p. 157–81.

40. Rakoczy-Trojanowska M, Bolibok-Brągoszewska H, Myśków B, Dzięgielewska M, Stojałowski S, Grądzielewska A, et al. Genetics and genomics of stress tolerance. In: Rabanus-Wallace MT, Stein N, editors. The Rye Genome, Compendium of Plant Genomes. Springer, Cham; 2021. p. 213–36.

41. Li G, Wang L, Yang J, He H, Jin H, Li X, et al. A high-quality genome assembly highlights rye genomic characteristics and agronomically important genes. Nat Genet. 2021;53:574–84.

42. Yuan Y, Gao M, Zhang M, Zheng H, Zhou X, Guo Y, et al. QTL Mapping for Phosphorus Efficiency and Morphological Traits at Seedling and Maturity Stages in Wheat. Front Plant Sci. 2017;8 April:1– 13.

43. Hawliczek A, Bolibok L, Tofil K, Borzęcka E, Jankowicz-Cieślak J, Gawroński P, et al. Deep sampling and pooled amplicon sequencing reveals hidden genic variation in heterogeneous rye accessions. BMC Genomics. 2020;21:845.

44. Hu B, Jin J, Guo AY, Zhang H, Luo J, Gao G. GSDS 2.0: An upgraded gene feature visualization server. Bioinformatics. 2015;31:1296–7.

45. Chen C, Chen H, Zhang Y, Thomas HR, Frank MH, He Y, et al. TBtools: An Integrative Toolkit Developed for Interactive Analyses of Big Biological Data. Mol Plant. 2020;13:1194–202.

46. Kumar S, Stecher G, Li M, Knyaz C, Tamura K. MEGA X: Molecular evolutionary genetics analysis across computing platforms. Mol Biol Evol. 2018;35:1547–9.

47. Zhan W, Cui L, Guo G, Zhang Y. Genome-wide identification and functional analysis of the TCP gene family in rye (Secale cereale L.). Gene. 2023;854 November 2022.

48. Madeira F, Madhusoodanan N, Lee J, Eusebi A, Niewielska A, Tivey ARN, et al. Using EMBL-EBI Services via Web Interface and Programmatically via Web Services. Curr Protoc. 2024;4:1–52.

49. Ye J, Coulouris G, Zaretskaya I, Cutcutache I, Rozen S, Madden TL. Primer-BLAST: A tool to design target-specific primers for polymerase chain reaction. BMC Bioinformatics. 2012;13:134.

50. Livak KJ, Schmittgen TD. Analysis of relative gene expression data using real-time quantitative PCR and the 2-ΔΔCT method. Methods. 2001;25:402–8.

51. R_Core_Team. R: A Language and Environment for Statistical Computing. Vienna, Austria; 2013.

52. Wickham H. ggplot2: Elegant Graphics for Data Analysis. New York: Springer New York; 2016.

53. Vaser R, Adusumalli S, Leng SN, Sikic M, Ng PC. SIFT missense predictions for genomes. Nat Protoc. 2016;11:1–9.

54. Tamura K, Stecher G, Kumar S. MEGA11: Molecular Evolutionary Genetics Analysis Version 11. Mol Biol Evol. 2021;38:3022–7.

55. Peakall R, Smouse PE. Genalex 6: genetic analysis in Excel. Population genetic software for teaching and research. Mol Ecol Notes. 2006;6:288–95.

56. Peakall R, Smouse PE. GenAlEx 6.5: genetic analysis in Excel. Population genetic software for teaching and research--an update. Bioinformatics. 2012;28:2537–9.

57. Saxena RK, Edwards D, Varshney RK. Structural variations in plant genomes. Briefings Funct Genomics Proteomics. 2014;13.

58. Ho SS, Urban AE, Mills RE. Structural variation in the sequencing era. Nat Rev Genet. 2020;21:171–89.

59. Yuan Y, Bayer PE, Batley J, Edwards D. Current status of structural variation studies in plants. Plant Biotechnol J. 2021;19:2153–63.

60. Zanini SF, Bayer PE, Wells R, Snowdon RJ, Batley J, Varshney RK, et al. Pangenomics in crop improvement—from coding structural variations to finding regulatory variants with pangenome graphs. Plant Genome. 2022;15:1–18.

61. Hattori Y, Nagai K, Furukawa S, Song XJ, Kawano R, Sakakibara H, et al. The ethylene response factors SNORKEL1 and SNORKEL2 allow rice to adapt to deep water. Nature. 2009;460:1026–30.

62. Heuer S, Lu X, Chin JH, Tanaka JP, Kanamori H, Matsumoto T, et al. Comparative sequence analyses of the major quantitative trait locus phosphorus uptake 1 (Pup1) reveal a complex genetic structure. Plant Biotechnol J. 2009;7:456–71.

63. Kong W, Wang Y, Zhang S, Yu J, Zhang X. Recent Advances in Assembly of Complex Plant Genomes. Genomics, Proteomics Bioinforma. 2023;21:427–39.

64. Bauer E, Schmutzer T, Barilar I, Mascher M, Gundlach H, Martis MM, et al. Towards a whole-genome sequence for rye (Secale cereale L.). Plant J. 2017;89:853–69.

65. Schatz MC, Witkowski J, McCombie WR. Current challenges in de novo plant genome sequencing and assembly. Genome Biol. 2012;13.

66. Fang Y, Jiang J, Hou X, Guo J, Li X, Zhao D, et al. Plant protein-coding gene families: Their origin and evolution. Front Plant Sci. 2022;13 September:1–15.

67. Rae AL, Cybinski DH, Jarmey JM, Smith FW. Characterization of two phosphate transporters from barley; evidence for diverse function and kinetic properties among members of the Pht1 family. Plant Mol Biol. 2003;53:27–36.

68. Dai C, Dai X, Qu H, Men Q, Liu J, Yu L, et al. The rice phosphate transporter OsPHT1;7 plays a dual role in phosphorus redistribution and anther development. Plant Physiol. 2022;188:2272–88.

69. Wang X, Wang Y, Piñeros MA, Wang Z, Wang W, Li C, et al. Phosphate transporters OsPHT1;9 and OsPHT1;10 are involved in phosphate uptake in rice. Plant, Cell Environ. 2014;37:1159–70.

70. Willmann M, Gerlach N, Buer B, Polatajko A, Nagy R, Koebke E, et al. Mycorrhizal phosphate uptake pathway in maize: Vital for growth and cob development on nutrient poor agricultural and greenhouse soils. Front Plant Sci. 2013;4 DEC:1–15.

71. Versaw WK, Harrison MJ. A chloroplast phosphate transporter, PHT2;1, influences allocation of phosphate within the plant and phosphate-starvation responses. Plant Cell. 2002;14:1751–66.

72. Guo C, Zhao X, Liu X, Zhang L, Gu J, Li X, et al. Function of wheat phosphate transporter gene TaPHT2;1 in Pi translocation and plant growth regulation under replete and limited Pi supply conditions. Planta. 2013;237:1163–78.

73. Wang J, Yang Y, Liao L, Xu J, Liang X, Liu W. Genome-wide identification and functional characterization of the phosphate transporter gene family in Sorghum. Biomolecules. 2019;9.

74. Smith FW, Mudge SR, Rae AL, Glassop D. Phosphate transport in plants. Plant Soil. 2003;248:71– 83.

75. Raghothama KG, Karthikeyan AS. Phosphate acquisition. Plant Soil. 2005;274:37–49.

76. Huang CY, Roessner U, Eickmeier I, Genc Y, Callahan DL, Shirley N, et al. Metabolite profiling reveals distinct changes in carbon and nitrogen metabolism in phosphate-deficient barley plants (Hordeum vulgare L.). Plant Cell Physiol. 2008;49:691–703.

77. Aziz T, Finnegan PM, Lambers H, Jost R. Organ-specific phosphorus-allocation patterns and transcript profiles linked to phosphorus efficiency in two contrasting wheat genotypes. Plant, Cell Environ. 2014;37:943–60.

78. Grün A, Buchner P, Broadley MR, Hawkesford MJ. Identification and expression profiling of Pht1 phosphate transporters in wheat in controlled environments and in the field. Plant Biol. 2018;20:374–89.

79. Anandan A, Parameswaran C, Mahender A, Nayak AK, Vellaikumar S, Balasubramaniasai C, et al. Trait variations and expression profiling of OsPHT1 gene family at the early growth-stages under phosphorus-limited conditions. Sci Rep. 2021;11:1–19.

80. Victor Roch G, Maharajan T, Ceasar SA, Ignacimuthu S. The Role of PHT1 Family Transporters in the Acquisition and Redistribution of Phosphorus in Plants. CRC Crit Rev Plant Sci. 2019;38:171–98.

81. Liu X li, Wang L, Wang X wen, Yan Y, Yang X li, Xie M yang, et al. Mutation of the chloroplast-localized phosphate transporter OsPHT2;1 reduces flavonoid accumulation and UV tolerance in rice. Plant J. 2020;102:53–67.

82. Lhamo D, Shao Q, Tang R, Luan S. Genome-wide analysis of the five phosphate transporter families in Camelina sativa and their expressions in response to low-P. Int J Mol Sci. 2020;21:1–17.

83. Wei X, Xu X, Fu Y, Yang X, Wu L, Tian P, et al. Effects of Soybean Phosphate Transporter Gene GmPHT2 on Pi Transport and Plant Growth under Limited Pi Supply Condition. Int J Mol Sci. 2023;24.

84. Ahmad I, Rawoof A, Islam K, Momo J, Ramchiary N. Identification and expression analysis of phosphate transporter genes and metabolites in response to phosphate stress in Capsicum annuum. Environ Exp Bot. 2021;190 July.

85. Jia F, Wan X, Zhu W, Sun D, Zheng C, Liu P, et al. Overexpression of mitochondrial phosphate transporter 3 severely hampers plant development through regulating mitochondrial function in Arabidopsis. PLoS One. 2015;10:1–14.

86. Kochian L V. Rooting for more phosphorus. Nature. 2012;488:466–77.

87. Hawliczek A, Borzęcka E, Tofil K, Alachiotis N, Bolibok L, Gawroński P, et al. Selective sweeps identification in distinct groups of cultivated rye (Secale cereale L.) germplasm provides potential candidate genes for crop improvement. BMC Plant Biol. 2023;23:323.

88. Monteiro F, Vidigal P, Barros AB, Monteiro A, Oliveira HR, Viegas W. Genetic distinctiveness of rye in situ accessions from Portugal unveils a new hotspot of unexplored genetic resources. Front Plant Sci. 2016;7.

89. Bolibok-Brągoszewska H, Targońska M, Bolibok L, Kilian A, Rakoczy-Trojanowska M. Genome-wide characterization of genetic diversity and population structure in Secale. BMC Plant Biol. 2014;14:184.

90. Li Y, Böck A, Haseneyer G, Korzun V, Wilde P, Schön C-C, et al. Association analysis of frost tolerance in rye using candidate genes and phenotypic data from controlled, semi-controlled, and field phenotyping platforms. BMC Plant Biol. 2011;11:146.

91. Varshney RK, Beier U, Khlestkina EK, Kota R, Korzun V, Graner A, et al. Single nucleotide polymorphisms in rye (Secale cereale L.): discovery, frequency, and applications for genome mapping and diversity studies. Theor Appl Genet. 2007;114:1105–16.

92. Bilton TP, Schofield MR, Black MA, Chagne D, Wilcox PL, Dodds KG. Accounting for Errors in Low Coverage High-Throughput. 2018;209 May:65–76.

93. Quiroz D, Lensink M, Kliebenstein DJ, Monroe JG. Causes of Mutation Rate Variability in Plant Genomes. Annu Rev Plant Biol. 2023;74:751–75.

94. Man J, Harrington TA, Lally K, Bartlett ME. Asymmetric Evolution of Protein Domains in the Leucine-Rich Repeat Receptor-Like Kinase Family of Plant Signaling Proteins. Mol Biol Evol. 2023;40:1–16.

95. Ohta T. Evolution of Gene Families. Brenner’s Encycl Genet Second Ed. 2013;259:563–5.

96. Manyuan Long. Evolution of novel genes Manyuan Long. 2001;:673–80.

97. Lynch M, Ackerman MS, Gout JF, Long H, Sung W, Thomas WK, et al. Genetic drift, selection and the evolution of the mutation rate. Nat Rev Genet. 2016;17:704–14.

98. Dwivedi SL, Ceccarelli S, Blair MW, Upadhyaya HD, Are AK, Ortiz R. Landrace Germplasm for Improving Yield and Abiotic Stress Adaptation. Trends Plant Sci. 2016;21:31–42.

99. Zhang H, Mittal N, Leamy LJ, Barazani O. Back into the wild — Apply untapped genetic diversity of wild relatives for crop improvement.2017; September 2016:5–24.

